# Intraoperative biomechanics of lumbar pedicle screw loosening following successful arthrodesis

**DOI:** 10.1101/057935

**Authors:** Hope B. Pearson, Christopher J. Dobbs, Eric Grantham, Glen L. Niebur, James L. Chappuis, Joel D. Boerckel

## Abstract

**Abstract:** Pedicle screw loosening has been implicated in recurrent back pain after lumbar spinal fusion, but the degree of loosening has not been systematically quantified in patients. Instrumentation removal is an option for patients with successful arthrodesis, but remains controversial. Here, we quantified pedicle screw loosening by measuring screw insertion and/or removal torque at high statistical power (β = 0.98) in N = 108 patients who experienced pain recurrence despite successful fusion after posterior instrumented lumbar fusion with anterior lumbar interbody fusion (L2-S1). Between implantation and removal, pedicle screw torque was reduced by 58%, indicating significant loosening over time. Loosening was greater in screws with evoked EMG threshold under 11 mA, indicative of screw misplacement. A theoretical stress analysis revealed increased local stresses at the screw interface in pedicles with decreased difference in pedicle thickness and screw diameter. Loosening was greatest in vertebrae at the extremities of the fused segments, but was significantly lower in segments with one level of fusion than in those with two or more.

**Clinical significance:** These data indicate that pedicle screws can loosen significantly in patients with recurrent back pain and warrant further research into methods to reduce the incidence of screw loosening and to understand the risks and potential benefits of instrumentation removal.

## Introduction

Chronic low back pain is the second most-common reason for visits to a physician in the United States, and interbody fusion is common in patients non-responsive to non-surgical options.^1^ In 2012, over 413,000 lumbar fusions were reported in the US, and demand for surgical treatment continues to rise.^2,3^ Improvements in surgical approaches,^4^ as well as fixation instrumentation^4,5^ and inductive bone formation agents^2,6,7^ have improved patient outcomes, reduced the frequency and extent of complications, and have collectively made lumbar spinal fusion a common procedure that has improved quality of life for millions of patients.

However, reoperation secondary to recurrent back pain has been reported in approximately 14-27% of patients,^8^ with complications including improper instrumentation placement, loss of fixation, fatigue and bending failure, dural tears, nerve root injury, infection, and pedicle screw loosening.^9,10^ While successful fusion is correlated with desirable clinical outcomes,^11^ recurrent pain can occur even in patients with solid arthrodesis, which may be associated with instrumentation loosening, requiring revision or removal.

Over the past ten years, posterior pedicle screw systems have increased in strength and rigidity,^12,13^ increasing pre-fusion stability and decreasing the incidence of pseudarthrosis.^13,14^ However, this has also increased the loads present at the bone-screw interface due to increased load-sharing,^15^ which may contribute to pedicle screw loosening.^16,17^ In addition, load sharing under compliant fixation can direct tissue differentiation, with moderate load transfer stimulating bone formation, but excessive motion inducing pseudarthrosis,^17^ potentially through inhibition of neovascularization.^18^ Surgical removal of pedicle screws has been much discussed,^10,19^ and remains controversial due to the associated risks of secondary surgery and potential for instability following removal.^20^ However, when unexplainable recurrent pain is severe, screw removal has been recommended,^21–24^ and may also reduce risks of metal toxicity and hypersensitivity.^25^

Importantly, the loosening of pedicle screws *in vivo* has not been thoroughly evaluated. Therefore, the aims of this study were to quantify pedicle screw loosening in patients with recurrent back pain following successful posterior instrumented lumbar fusion (PILF) surgery and to assess patient pain perception prior to and following instrumentation removal.

## Methods

This is a retrospective cohort study with level of evidence of Level II.

### Patients

A total of 108 patients (75 male, 33 female), 29-78 years of age (mean 47, SD 11), were voluntarily enrolled following exclusion criteria according to STROBE (strengthening the reporting of observational studies in epidemiology) guidelines (Figure 1A). A single surgeon (JLC) performed all procedures. Each patient had been diagnosed with degenerative spinal stenosis with associated instability, degenerative spondylolisthesis, or annular tears and discogenic pain that had failed conservative treatment for at least one year. All subjects gave informed consent to participate, and the study was approved by Sterling Institutional Review Board, Atlanta, GA (#5187). Patients received 2- to 5-level spinal fusion of lumbar vertebrae between L2 and S1 (Figure 1B).

Only patients with successful arthrodesis were selected for subsequent evaluation. Patients who returned with complaints of recurrent pain that could not be explained by alternative diagnoses such as insufficient fusion or pseudarthrosis, infection, instrumentation failure, or soft tissue injury, and who elected both instrumentation removal and informed inclusion in the study were selected for analysis. To verify that recurrent pain was instrumentation-associated, anterior and posteriolateral computed tomography (CT) scans were assessed to rule out pseudarthrosis, adjacent segment disease, and sagittal imbalance. Further, following a radiopaque dye injection (10cc Amipaque) under C-arm guidance to provide X-ray contrast, a peri-implant “hardware block” was performed by Marcaine injection (1%, 20cc), and ambulatory patient pain was evaluated. Only patients with successful arthrodesis whose pain could be attributed to the instrumentation by hardware block were then selected for instrumentation removal in a second surgery. Measurements at both insertion and removal were taken in thirty-seven patients (12 male, 25 female) for paired assessment. Measurements at insertion only were assessed in thirty-five patients (32 male, 3 female), and measurements at removal only were taken in thirty-six patients (31 male, 5 female).

Removal torque measurement was assessed at an average ± SD of 266 ± 73 days post-implantation.

**Figure 1.**
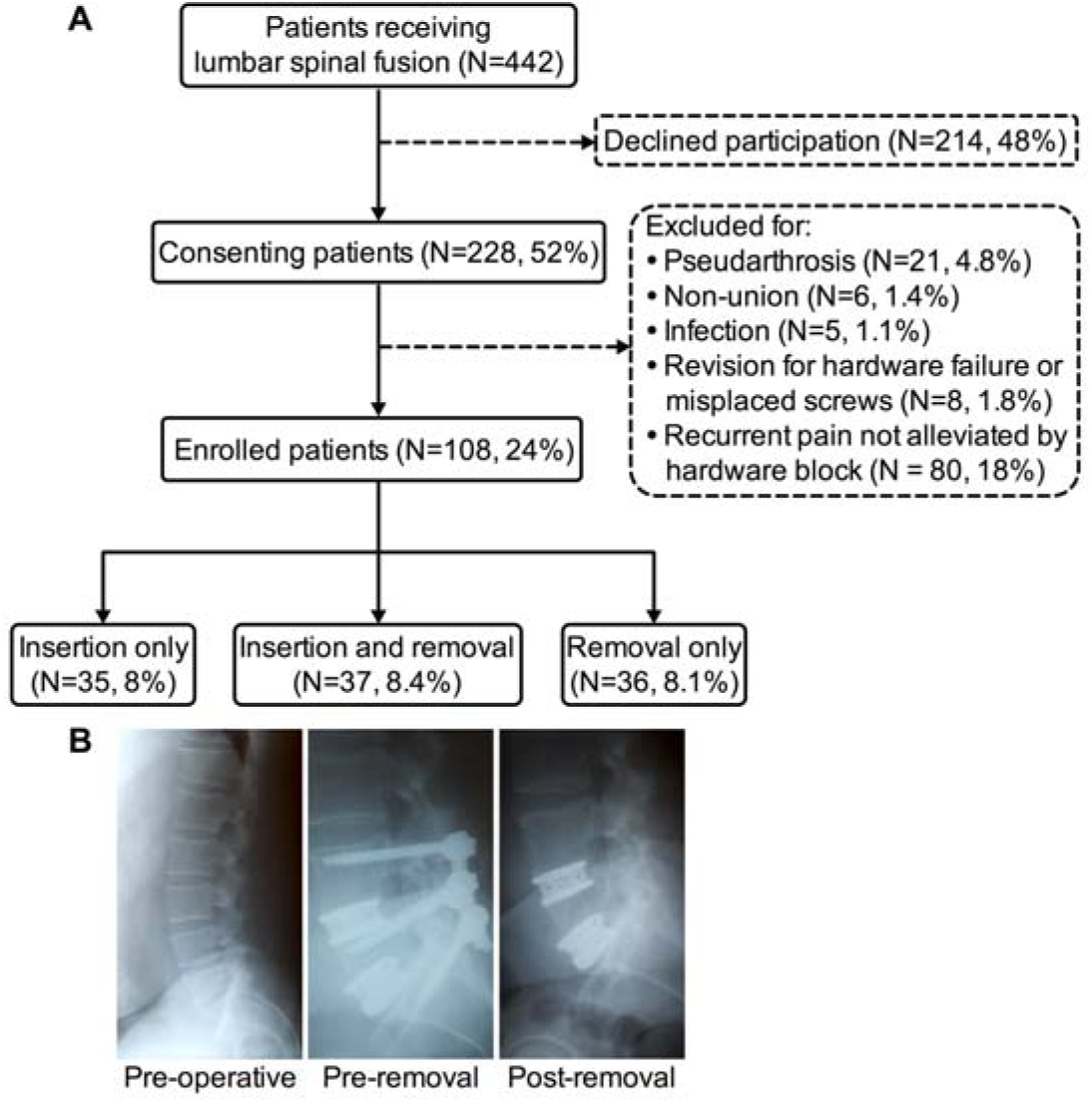
Patient selection and pedicle screw removal. (A) STROBE flowchart of subject inclusion and exclusion. (B) Representative X-ray images of pedicle screw removal following successful arthrodesis at pre-operative, post-fusion, and post-removal time points.

### Surgical approach

Patients received posterior instrumented lumbar fusion (PILF) combined with anterior lumbar interbody fusion (ALIF) to maximize segment stability (Figure 2A). Briefly, patients were placed prone on a Jackson operating table, prepped, and draped in usual sterile fashion. After open exposure of the pedicles, pedicle holes were tapped and titanium pedicle screws (TSRH-3D®) were inserted under C-arm guidance based on the standard progression of coronal angle (5° at L1, progressing 5° at each successive caudal level). In the saggital plane, screws were inserted level with the pedicle and disc space without caudal/cephalad angling. Screw diameters were chosen according to manufacturer recommendations, and lengths were selected to achieve 60-80% engagement in the vertebral body upon full insertion. Holes were tapped 1mm diameter smaller than the screw size (i.e., undertapped), yielding a screw-hole interference fit, *δ_i_* = *r_s_* − *r_h_* = 0.5 mm, where *r_s_* is the radius of the pedicle screw, and *r_h_* is the radius of the tapped hole prior to screw insertion.^26^

The decision to proceed with instrumentation removal in consenting patients was made upon verification of successful arthrodesis by computed tomography scan in both anterior and posterolateral views and nullification of alternative explanations. A total of 80 patients with recurrent pain did not receive hardware removal due to inability to identify the hardware as cause of recurrent pain (Figure 1A). Selected patients were returned to the operating room no sooner than 6 months after insertion and up to 3 years (mean ± SD 266 ± 73 days) after the initial procedure. Instrumentation was exposed in routine fashion, and side connectors and rods were removed. Screw complications were not observed prior to removal. Beam-hardening artifacts limited our ability to assess peri-implant radiodensity by computed tomography on post-implantation and pre-removal scans.

### Pedicle screw insertion and removal torque

Pedicle screw insertion (N = 467) and removal (N = 477) torques were measured using a sterilizeable, manual mechanical torque wrench (Mountz, San Jose, CA), with N = 139 paired insertion and removal measurements in the same patients. To ensure changes in torque between insertion and removal surgeries did not occur due to hole tapping by the insertion screw, pedicle screw back-out torque (at the time of insertion) was also measured (N = 204).

### Evoked EMG stimulus threshold

Paired evoked electromyographic (EMG) stimulus thresholds through the screw hole^27^ were measured in N = 98 screws. Non-paired measurements at insertion (N = 394) and removal (N = 430) were also taken.

### Pedicle and screw measurements

Vertebra level, screw type, diameter, and length were recorded for each measurement. Pedicle thickness was measured from pre-operative digital CT scans in the transverse plane (Figure 4A), and pedicle-screw diameter difference, defined as pedicle thickness minus screw diameter was computed.

### Stress analysis

A theoretical stress analysis of monotonic cantilever loading in transverse compression and bending and press-fit by screw insertion was performed to estimate the stresses in peri-implant pedicle bone with or without loosening of the screw. The trabecular bone of the pedicle was assumed to have an effective modulus of 42.8 MPa and a Poisson ratio of 0.25, while the cortical bone was assigned a modulus of 21 GPa and a Poisson ratio of 0.37 (Appendix Table A1).^28–31^ The modulus of the titanium screws was assumed to be 180 MPa. Loading of the spine causes cantilevered loads and moments to be transmitted through the instrumentation, resulting in both a transverse load and moment to be distributed to the surrounding trabecular bone along the embedded length of the screw. The stresses induced by this cantilever loading can be approximated by beam-on-elastic-foundation theory,^31^ assuming the screw and cortical bone act as co-axial deformable beams joined by an elastic foundation of trabecular bone, assumed here to be homogeneous. See Appendix 1 for the associated differential equations, boundary conditions, and solution approach. We evaluated the stress distribution in the trabecular bone along the length of the pedicle screw and computed the maximal stress, *σ_r_cantilever__*, which occurred at the screw insertion point (*Z* = *L*; labeled as point A in Figure 5A). Since the flexural rigidity of the trabecular bone is substantially lower than either the cortical bone or titanium screw, the axial stresses in the elastic trabecular layer due to bending can be neglected. Local stresses were assessed under physiologic loading conditions of standing, extension, and flexion^32,33^ with assumptions of either loosened or bonded screws, as described by Huiskes.^31^ Next, a press-fit analysis based on linear elastic thick-walled pressure vessel theory was used to determine the “locked-in” stress induced by the insertion of the screw for the given screw-hole interference fit (*δ_i_* = 0.5*mm*. See Appendix 2 for associated differential equations, boundary conditions, and solution approach. To determine the composite stress state as a superposition of the residual stresses from the combined cantilevered loading and interference fit, the maximum von Mises stress in the pedicle at the screw interface, σ_H_, was defined as:

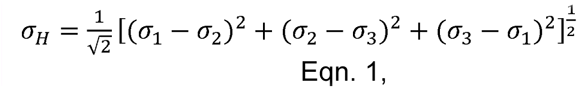

Where *σ*_1_ = *σ_r_interference__* + *σ_r_cantilever__*, *σ*_2_ = *σ_θ_interference__*, and *σ*_3_ ≈ 0.

### Visual analogue scale assessment of patient pain

Finally, to evaluate patient perception of lumbar pain, patients (N = 31 complete responders) were self-assessed using the visual analog scale (VAS; 0-100 mm) prior to fusion surgery, 1 month after insertion, prior to removal, 0-3 months after removal, and 3-6 months after removal. Self-reporting of the degree of analgesic medicine use and activity were also assessed using the same VAS approach.

### Statistical analysis

Differences between groups were evaluated by two-tailed Student’s t-test or analysis of variance (ANOVA) followed by Tukey’s post-hoc test for single and multiple comparisons, respectively, when assumptions of normality and homoscedasticity were met by D’Agostino-Pearson omnibus normality and F tests, respectively. Otherwise, non-parametric Mann-Whitney and Kruskal-Wallis followed by Dunn’s multiple comparison tests were used. P-values less than 0.05 were considered significant. For paired measurements, differences were analyzed by paired Student’s t-test or Wilcoxon’s matched-pairs signed rank test. Correlations between groups were assessed by linear regression using Pearson’s coefficient of determination (R^2^).

**Figure 2.**
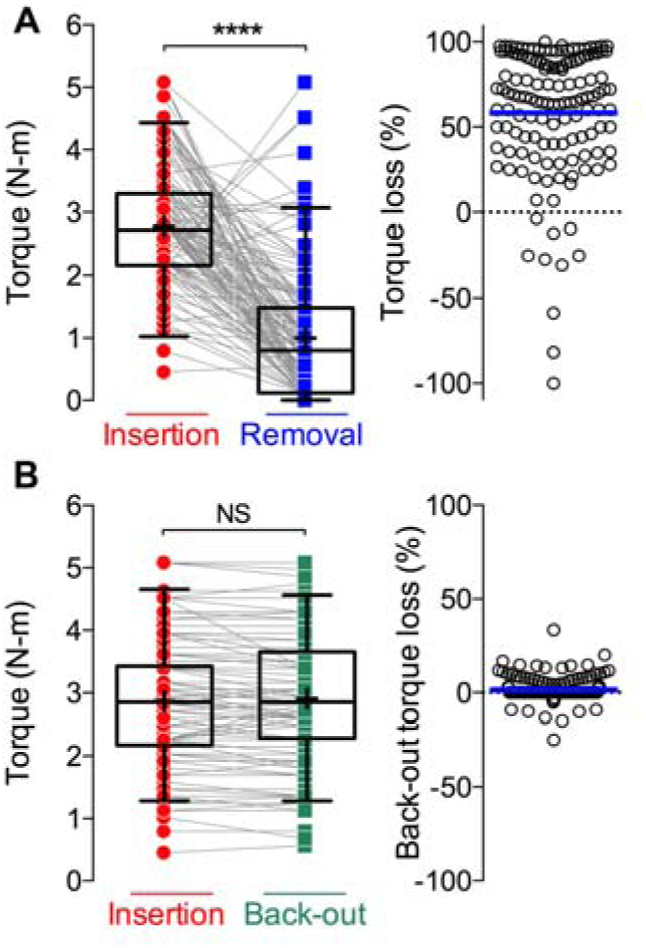
Measurement of pedicle screw loosening by insertion and removal torque. (A) Insertion and removal torque with paired samples connected by gray lines (N = 139 pairs), and scattergram of percent torque loss for those same pairs. (B) Insertion and immediate back-out torque (N = 204 pairs). Box plots show median line and 25^th^, and 75^th^ percentiles, with whiskers at 5^th^ and 95^th^ percentiles, respectively. Mean values indicated by + symbol. Scattergrams show mean values indicated by blue line. Statistical differences assessed by paired two-tailed Student’s t-test. **** p < 0.0001, NS not significant.

## Results

### Insertion and removal torque

Paired insertion and removal torque measurements (N = 139 pairs) revealed a significant 58.1% lower removal torque compared to insertion of those same screws (p < 0.0001), with only 9% of screws loosening by less than 15% (Figure 2A). To ensure that this reduction in torque between insertion and removal was not simply due to tapping of the screw hole during insertion of the screw, the single-rotation back-out torque was also evaluated at the time of insertion. The immediate torque loss between insertion and back-out was 1.4 ± 5.6% (Figure 2B; p = 0.68, β > 0.98). Samples were clustered at a mean percent change of 0% (Figure 2B, right). Non-paired insertion (N = 467) and removal torques (N = 477) were also evaluated to determine whether a similar difference in torque would also be apparent in non-paired samples, and revealed a similar 68% difference in torque between insertion and removal (p < 0.0001, Supplemental Figure 1A).

### Evoked EMG stimulus threshold

For paired samples, the difference in stimulus A difference in stimulus threshold at the screw hole was not measurable between insertion and removal (p > 0.05, β = 0.89; Figure 3A). However, torque loss in samples with less than 11 mA threshold at insertion, which has been identified as an indicator of screw placement and pedicle cortex violation,^27^ was significantly greater than those with insertion threshold above 11 mA (p < 0.05, Figure 3B). For unpaired measurements, the stimulus threshold at screw removal was 15% lower than the stimulus threshold at insertion (p < 0.0001, N = 427, 393, respectively; Supplemental Figure 1B).

**Figure 3.**
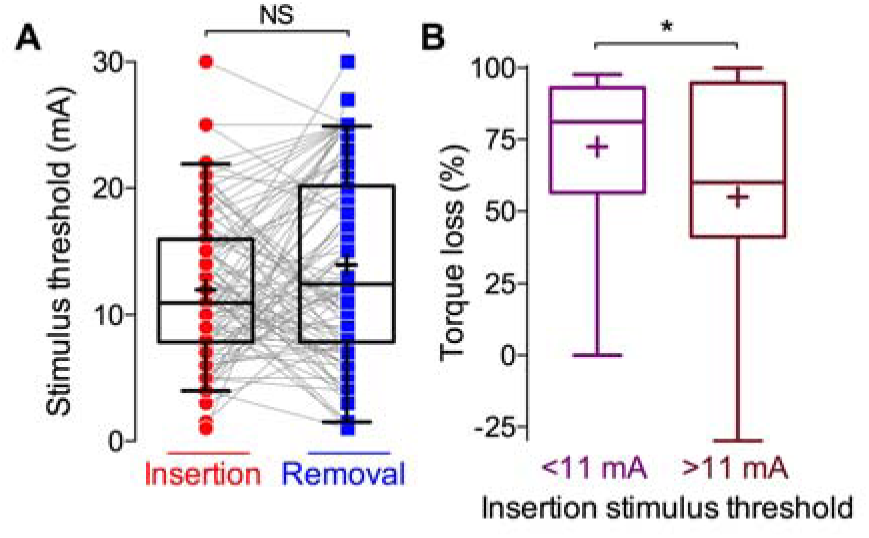
Evoked EMG stimulus threshold at insertion and removal. (A) Insertion and removal stimulus threshold with paired samples connected by gray lines (N = 98 pairs). (B) Percent torque loss for paired samples with insertion stimulus threshold less than (purple) or greater than (brown) the 11 mA cutoff indicative of pedicle screw placement quality.^27^ Box plots show median line and 25^th^, and 75^th^ percentiles, with whiskers at 5^th^ and 95^th^ percentiles, respectively. Mean values indicated by + symbol. Statistical differences assessed by paired (A) and unpaired (B) two-tailed Student’s t-test. * p < 0.05, NS not significant.

**Figure 4.**
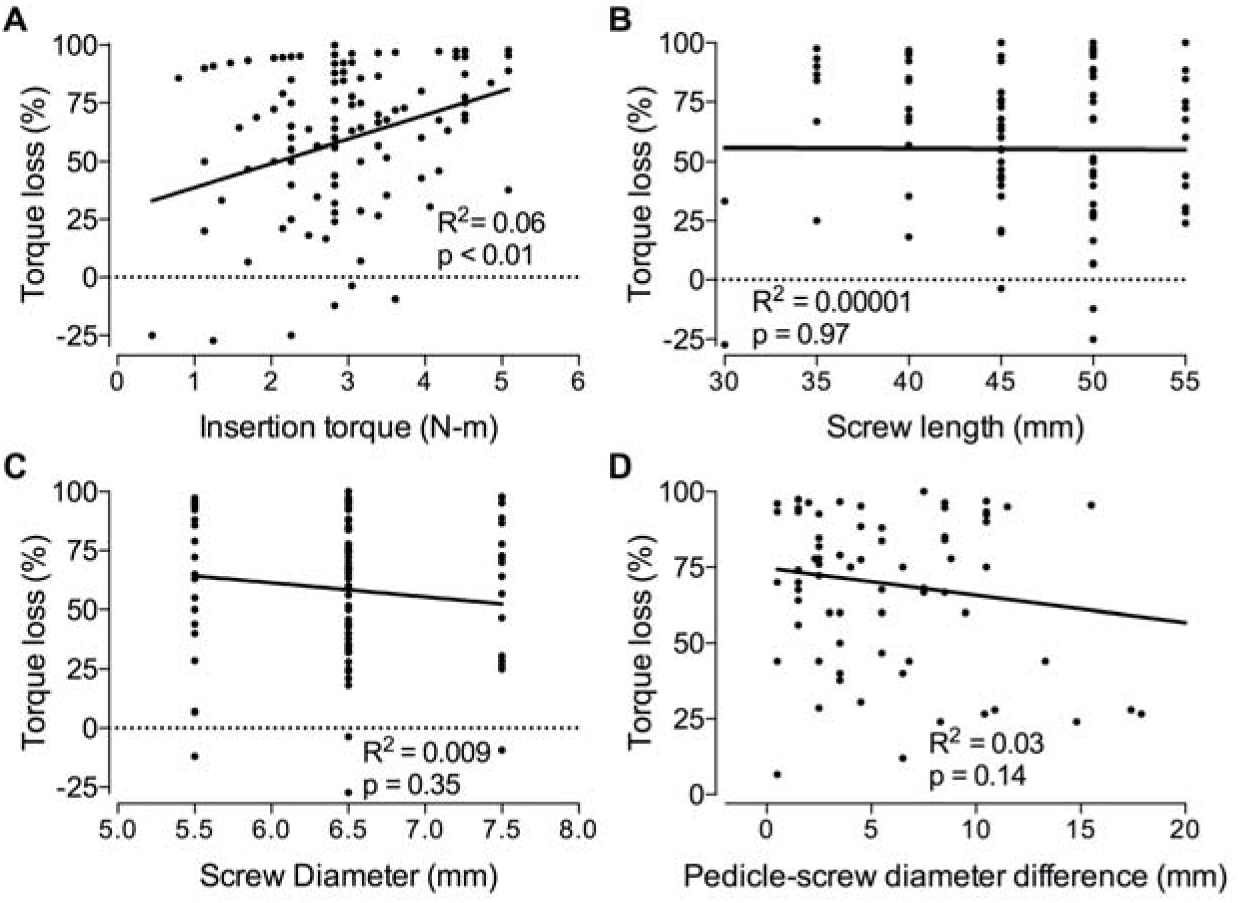
Linear regression analysis of pedicle screw loosening for intraoperative measurements. Correlation of percent torque loss with: (A) insertion torque (N = 139), (B) screw length (N = 114), (C) screw diameter (N = 139), (D) pedicle-screw diameter difference (N = 106).

**Figure 5.**
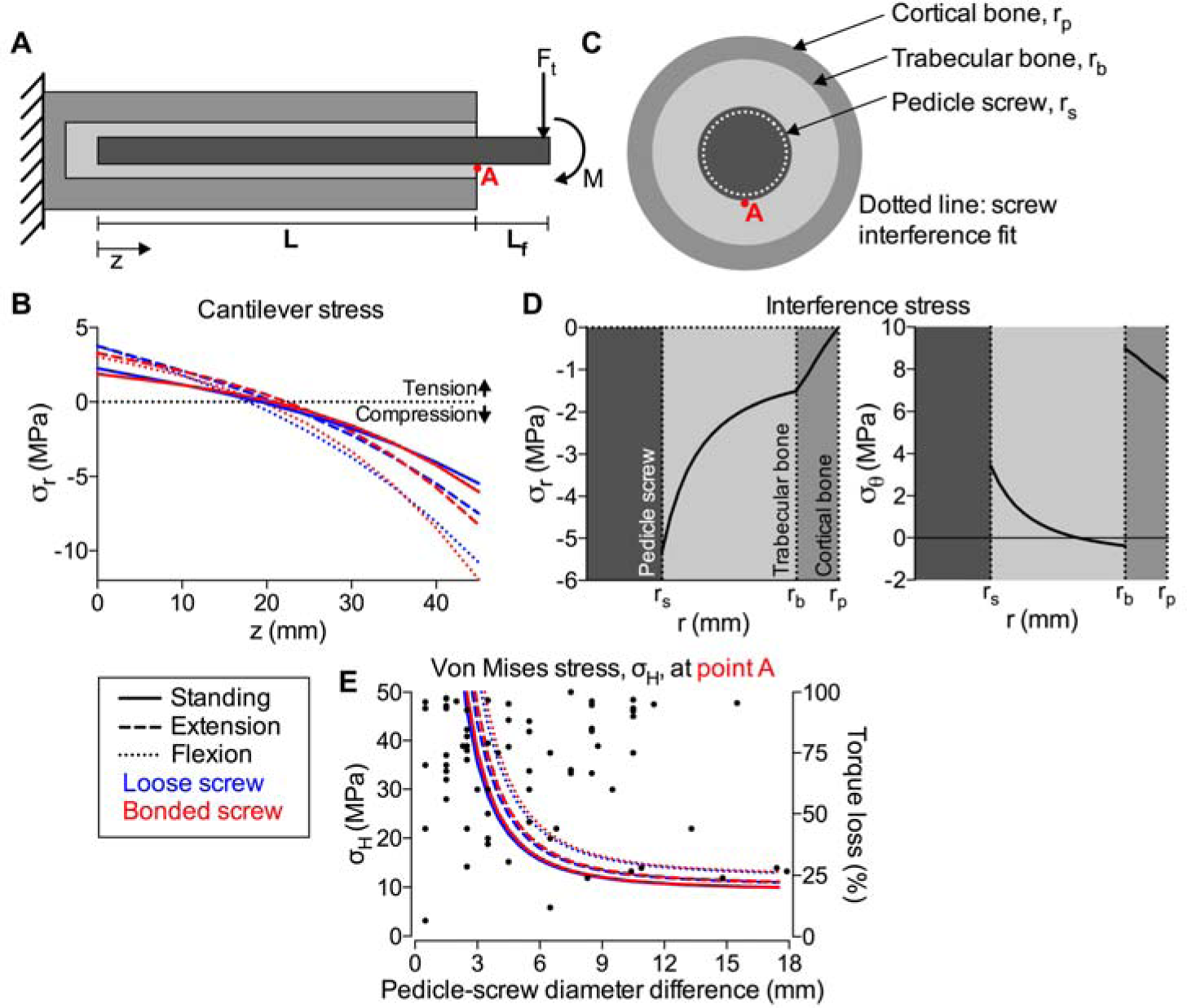
Analysis of stresses induced by cantilevered instrumentation loading (F_t_, M) and screw interference fit. (A) Schematic of the screw and pedicle approximated as co-axial beams joined by an elastic foundation of trabecular bone. (B) Stress distribution along the screw under physiologic loading conditions for standing, extension, and flexion of the spine assuming either the screw is either loose (blue) or perfectly bonded (red). (C) Schematic of interference fit. (D) Radial and circumferential stresses induced by screw insertion assuming an interference fit of *δ_i_* = *r_s_* − *r_h_* = 0.5 mm. (E) Von Mises stress at the location of maximal stress (labeled point A) as a function of pedicle-screw diameter difference superimposed on percent torque loss as measured in patients.

### Correlation analysis

Correlation analyses were performed to determine whether insertion torque, screw geometry, or relative fit in the pedicle could predict loosening. Insertion torque correlated with screw loosening (p < 0.01), but with little predictive power (R^2^ = 6%; Figure 4A). Neither screw diameter nor length^34^ predicted the extent of loosening, even in long screws with increased vertebral body penetration (Figure 4B, C). Torque loss was weakly correlated (p = 0.14, R^2^ = 3%) with pedicle screw-diameter difference, defined as the difference between pedicle thickness and screw diameter, as a measure of geometric pedicle integrity post-insertion (Figure 4D).^35^ EMG stimulus threshold did not linearly correlate with loosening either at insertion or at removal (Supplemental Figure 2).

### Stress analysis

Analytical models were used to estimate the stress distribution in the peri-implant bone along the length of the screw for cantilevered screw loads and moments using reported values for physiologic static loading conditions under standing, extension, and flexion poses^32,33^ (Figure 5A, B). Screw-hole interference stresses were calculated to approximate the press-fit-induced radial and circumferential stresses in the trabecular and cortical bone as a function of radial position (Figure 5C, D). The location of maximal von Mises stress occurred at the insertion point, immediately adjacent to the screw (indicated by Point A in Figure 5A, C). Under the applied loading conditions, the local von Mises stress decreased with increasing pedicle-screw diameter difference, shown overlaid with measured torque loss (Figure 5E). The degree of pedicle fill substantially influenced local stress values, in a manner dependent on both applied loads and interference stresses.

### Loosening by vertebra level

The percent torque loss by vertebra level in paired samples was evaluated at five levels of vertebrae from L2-S1 (Figure 6A; pairings showing adjacent segments within the same patient illustrated by connecting lines). The greatest degree of loosening was observed in L2 and S1, and the least in L4 (p < 0.05 by ANOVA with Tukey’s post-hoc test). To determine whether this was inherent to the vertebral level or whether the fused segments at the ends of the fusion exhibit more loosening, we compared the torque loss in the superior and inferior extremes of the fused segments vs. internal segments, and found greater loosening in vertebrae at the edges of the fusion segment (p < 0.05; Figure 6B). No differences were found between the superior-most vs. inferior-most segments. Next, we evaluated the mean torque loss as a function of the number of fused vertebrae (Figure 6C). While all levels exhibited significant loosening, screws from segments with a single level of fusion exhibited measurably less loosening compared to those with multiple levels of fusion (p < 0.05). Similar trends were observed for non-paired samples (Supplemental Figure 3).

### Patient pain perception

While pain perception in these patients was significantly reduced by 19% within 1 month after surgery, pain was recurrent, and returned to 89% of pre-surgery levels, causing patients to return for instrumentation removal surgery (Figure 7). Following removal, mean VAS pain scores returned to post-insertion levels, but differences with pre-removal pain were not statistically significant. Three to nine months after removal surgery, mean VAS pain scores remained at 27% lower than the pre-fusion levels. No differences were found in patient-reported use of analgesics or physical activity between any time points (Supplemental Figure 4).

**Figure 6.**
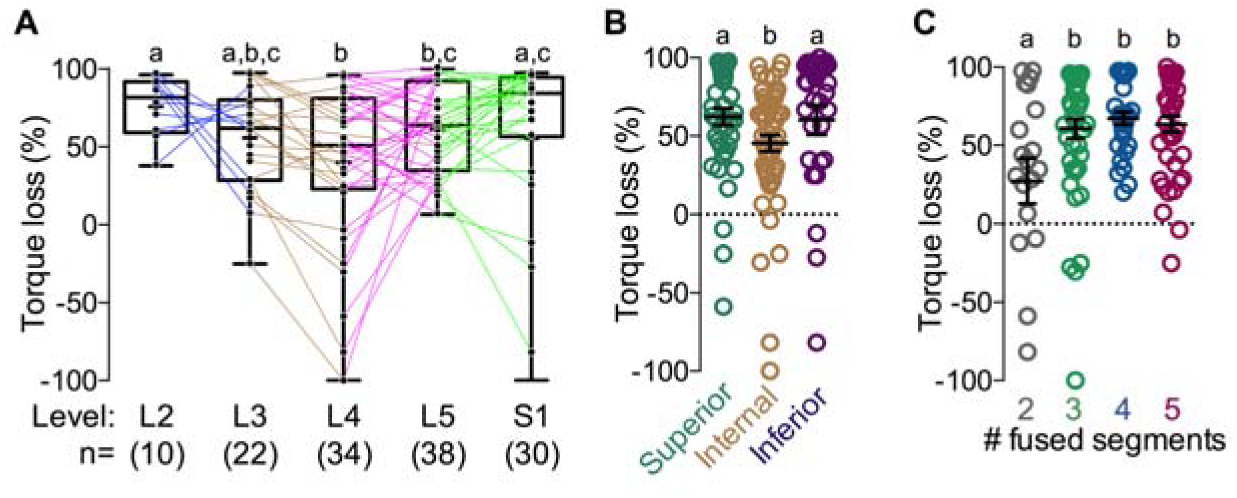
Pedicle screw loosening by vertebra level. (A) Percent torque loss for vertebra levels L2-S1 for paired measurements (N = 139). Box plots show median line and 25^th^, and 75^th^ percentiles, with whiskers at 5^th^ and 95^th^ percentiles, respectively. Mean values indicated by + symbol. (B) Torque loss in internal segments vs. those at the extremes of the fusion segment (superior/inferior), shown with mean ± s.e.m. (C) Torque loss as a function of number of levels of fusion, shown with mean ± s.e.m. Significance indicator letters shared in common between groups indicate no significant difference in pairwise comparisons. * indicates significant differences between superior/inferior and internal groups.

**Figure 7.**
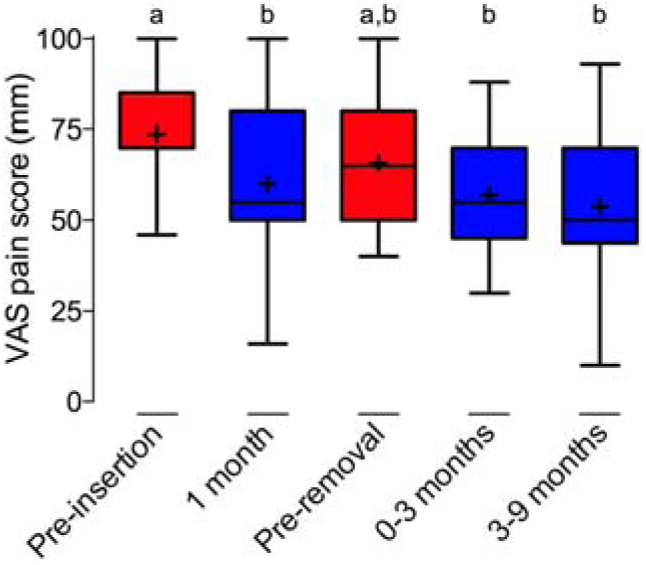
Patient assessment of recurrent pain. Visual Analog Scale (VAS; 0-100mm) assessment of pain (N = 31). Box plots show median line and 25^th^, and 75^th^ percentiles, with whiskers at 5^th^ and 95^th^ percentiles, respectively. Mean values indicated by + symbol. Significance indicator letters shared in common indicate no significant difference in pairwise comparisons.

## Discussion

Taken together, these data indicate that pedicle screws can loosen over time by an average of 58%, even after successful arthrodesis, in patients with recurrent back pain. This is significant because, while screw loosening on removal has been observed previously in a small pilot study,^36^ this is the first demonstration of the prevalence and degree of this problem. Sanden et al. evaluated pedicle screw loosening in eight patients, observing a 61% reduction in torque, which is highly consistent with our observations.^36^ They found that insertion torque correlated with removal torque in six screws, but the remaining measurements exhibited no discernable correlation. Consistent with this observation, our data suggest that only 6% of the variation in loosening can be predicted by insertion torque. Additionally, our data implicate screw placement, pedicle-screw diameter difference, fusion level, and number of fused vertebrae as important biomechanical factors in screw loosening.

Certainly, measures that could predict loosening *a priori* would have significant clinical utility, though this is challenging due to the multifactorial nature of screw loosening. Other studies have also evaluated insertion torque as a potential predictor, with some proposing this measure as a better predictor than bone mineral density,^37–39^ but its correlation with linear pullout strength in cadaveric studies is weak (R^2^=0.04-0.4).^40,41^ Consistent with these observations, we found that insertion torque correlated significantly with torque loss, but the predictive power was poor (R^2^ = 6% for N = 139pairs), indicating that intraoperative measurement of pedicle screw insertion torque is not uniquely sufficient to predict the degree of loosening.

Evoked EMG stimulus threshold is frequently used to detect screw placement and pedicle cortex violation, and could also be indicative of osteopenia.^27,42^ In the present study, intraoperative EMG measurements indicated that screws with EMG thresholds less than 11mA, indicative of poorly-seated screws or screws in osteopenic bone, have a greater likelihood of loosening; however, EMG stimulus threshold could not linearly predict the degree of loosening. These observations suggest that further research is warranted to decouple the interactions between local mineral density, screw placement and angle, and pedicle screw loosening. The evoked EMG stimulus threshold is commonly used to assess pedicle wall violation and nerve root exposure,^27^ so screw loosening may therefore induce late-onset nociceptive pain through stimulation of mechanosensitive nerve fibers upon progressive degradation of the peri-implant bone matrix.

The degree of loosening was vertebra level-dependent in a biphasic manner, with the least amount of loosening in the middle of the range of operated segments, and increased loosening in vertebrae at the extremities of the fused segments; however, screws in patients with a single level of fusion exhibited measurably lower torque loss than those with fusion of three or more vertebrae. Together, these data implicate biomechanical stability as an important factor in pedicle screw loosening.

The mechanisms that contribute to pedicle screw loosening are certain to be multifactorial, with both mechanical and biological components. One possibility is instrumentation-induced stress shielding and subsequent disuse osteopenia. A decrease in bone mineral density under stiff fixation has been seen in both canine^43,44^ and human^45^ implanted fixation devices as a result of stress shielding. Conversely, transfer of moderate levels of mechanical stimuli through compliant fixation can enhance BMP-2-induced bone formation in large defects, but excessive motion can induce pseudarthrosis and non-union^17,18,46^ Thus, the optimal rigidity of spinal fixation instrumentation remains a subject of ongoing investigation. Another likely contributor is local plastic deformation at the bone-screw interface caused by the insertion and cyclic loading of the pedicle screws, resulting in screw toggling and loosening. Pedicle screw insertion also induces residual stresses and stress concentrations that superimpose with cyclic loads to contribute to local tissue failure. These stresses are highest at the cap-rod-screw interface,^47,48^ as we demonstrate using beams-on-elastic-foundation and thick-wall pressure vessel theory. Reported yield stresses for spinal trabecular bone range from 3 to 96 MPa^49,50^. Using the von Mises stress as a measure of failure propensity, we calculated local stresses at the screw entry point of 50 MPa at median pedicle/instrumentation geometries, suggesting that local stresses may contribute to loosening through load-induced toggling. These values are consistent with prior reports of static vertebral von Mises stresses up to 100 MPa at the pedicle body junction. Incorporation of pedicle screw loosening into the model revealed increased stresses at the screw insertion point following screw loosening, which would positively feed back to promote further loosening consistent with a prior finite element comparisons of stresses around loosened and bonded pedicle screws.^47^ A recent hybrid multibody/finite element study by Fradet et al.^50^ evaluated the stresses induced by simulated intraoperative scoliosis correction maneuvers as well as failure loading of the instrumented spine, without fusion. They identified surgical maneuvers induced stresses of 40 - 100 MPa at the screw entry point.^
50^ Notably, their model, which featured L1-L3 fusion, identified maximal screw interface stresses in the proximal-most segment, L1, consistent with the loosening trends observed here. Together, these data suggest that both biological and mechanical factors influence the etiology of pedicle screw loosening, and further research is needed to elucidate the mechanisms underlying the initiation and progression of screw loosening as well as clinical implications for surgical approach and post-operative pain management.

This model did not account for bone remodeling induced by local damage or biological responses to the implant, which can also change and progress over time.^51^ High static stresses, combined with cyclic fatigue loading, may drive local pedicle failure and cause loosening over time. Screw trajectory also influences the biomechanical environment^52^ and its impact on loosening requires further research. Our observations together suggest that increasing screw size relative to either pedicle thickness or initial hole diameter may impair long-term stability as a result of local stress concentration. Future studies will be required to evaluate whether patients with low pedicle thickness exhibit increased likelihood of loosening, requiring additional observation or screw augmentation.

While the full clinical implications of pedicle screw loosening in balance with the potential negative sequelae of spinal instrumentation removal will require further study, these data suggest that pedicle screw loosening is prevalent in patients with recurrent pain, despite successful arthrodesis. This study was unable to determine whether screw loosening is present in patients without recurrent pain since unwarranted instrumentation removal for purely observational purposes would not be in the best interests of the patients. However, our observations of patient pain perception are in agreement with several studies, suggesting that pedicle instrumentation removal following successful arthrodesis may relieve recurrent pain,^21–24^ potentially caused by pedicle screw loosening. Further research will be necessary to fully evaluate clinical and pain outcomes and to establish pedicle screw loosening as a causative factor.

## SUPPLEMENTAL DATA

**Supplemental Figure 1.**
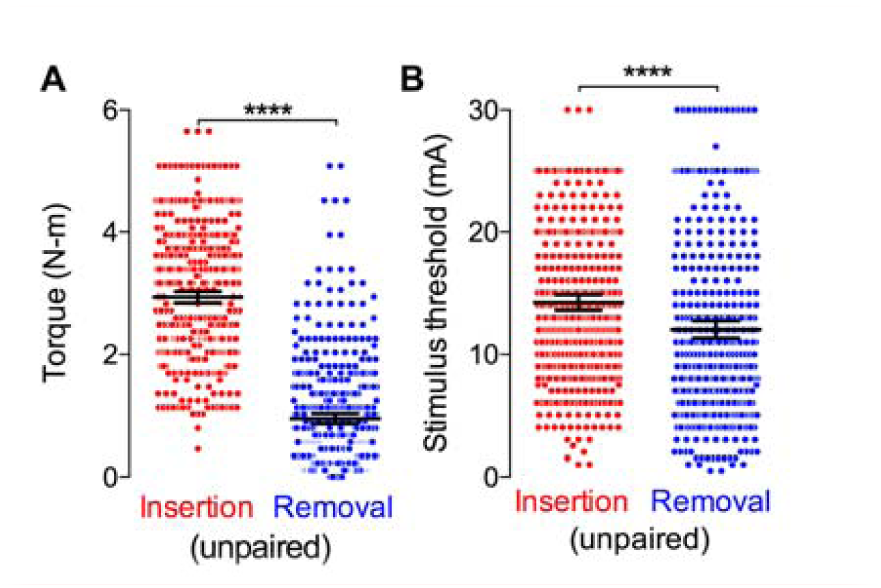
Insertion/removal torque and evoked EMG stimulus threshold in unpaired samples. (A) Insertion and removal torque (N = 467 and 477 for insertion and removal, respectively). (B) stimulus threshold (N = 394 and 430 for insertion and removal, respectively).

**Supplemental Figure 2.**
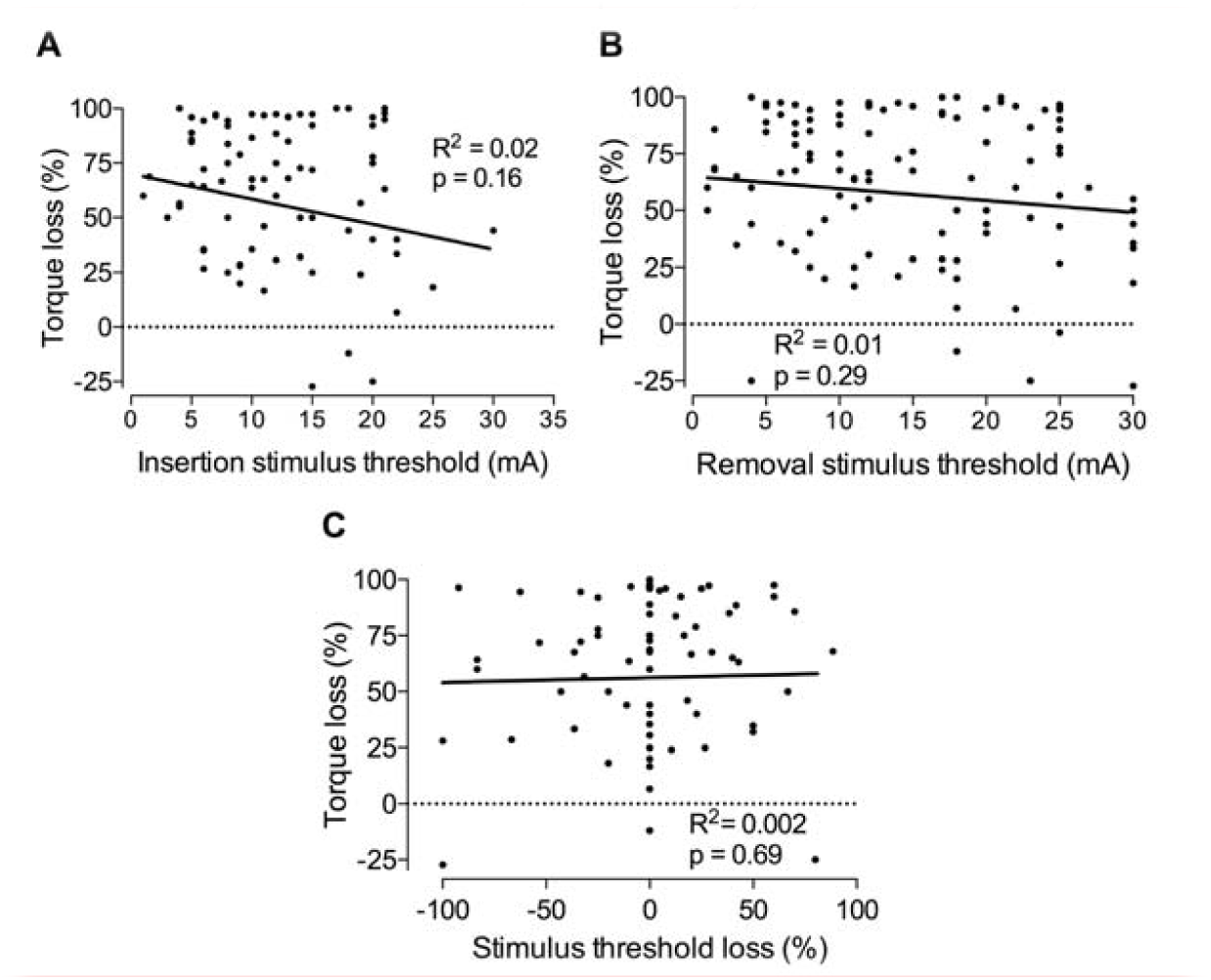
Linear regression analysis of pedicle screw loosening for intraoperative stimulus threshold measurements. Correlation of percent torque loss with (A) insertion stimulus threshold (N = 98 pairs), (B) removal stimulus threshold (N = 114 pairs), and (C) percent stimulus threshold loss for paired samples (N = 84 pairs). p > 0.05 indicates the slope of the regression line is not significantly different from zero.

**Supplemental Figure 3.**
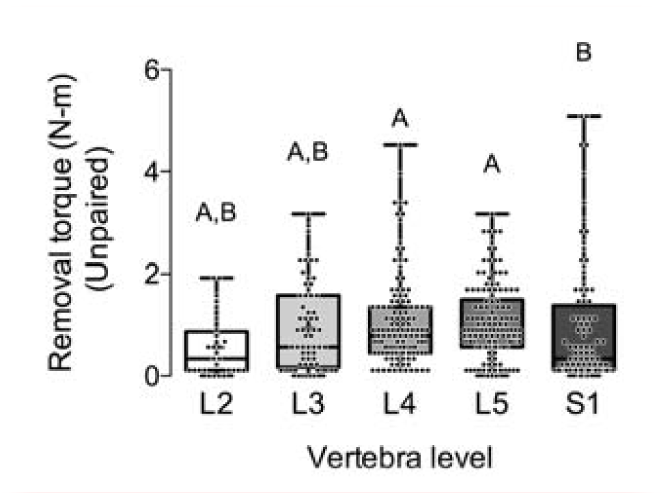
Pedicle screw removal torque by vertebra level. Unpaired removal torques for vertebra levels L2-S1 (N = 430). Box plots show median line and 25^th^, and 75^th^ percentiles, with whiskers at 5^th^ and 95^th^ percentiles, respectively. Mean values indicated by + symbol. Significance indicator letters shared in common between or among vertebra level groups indicate no significant difference.

**Supplemental Figure 4.**
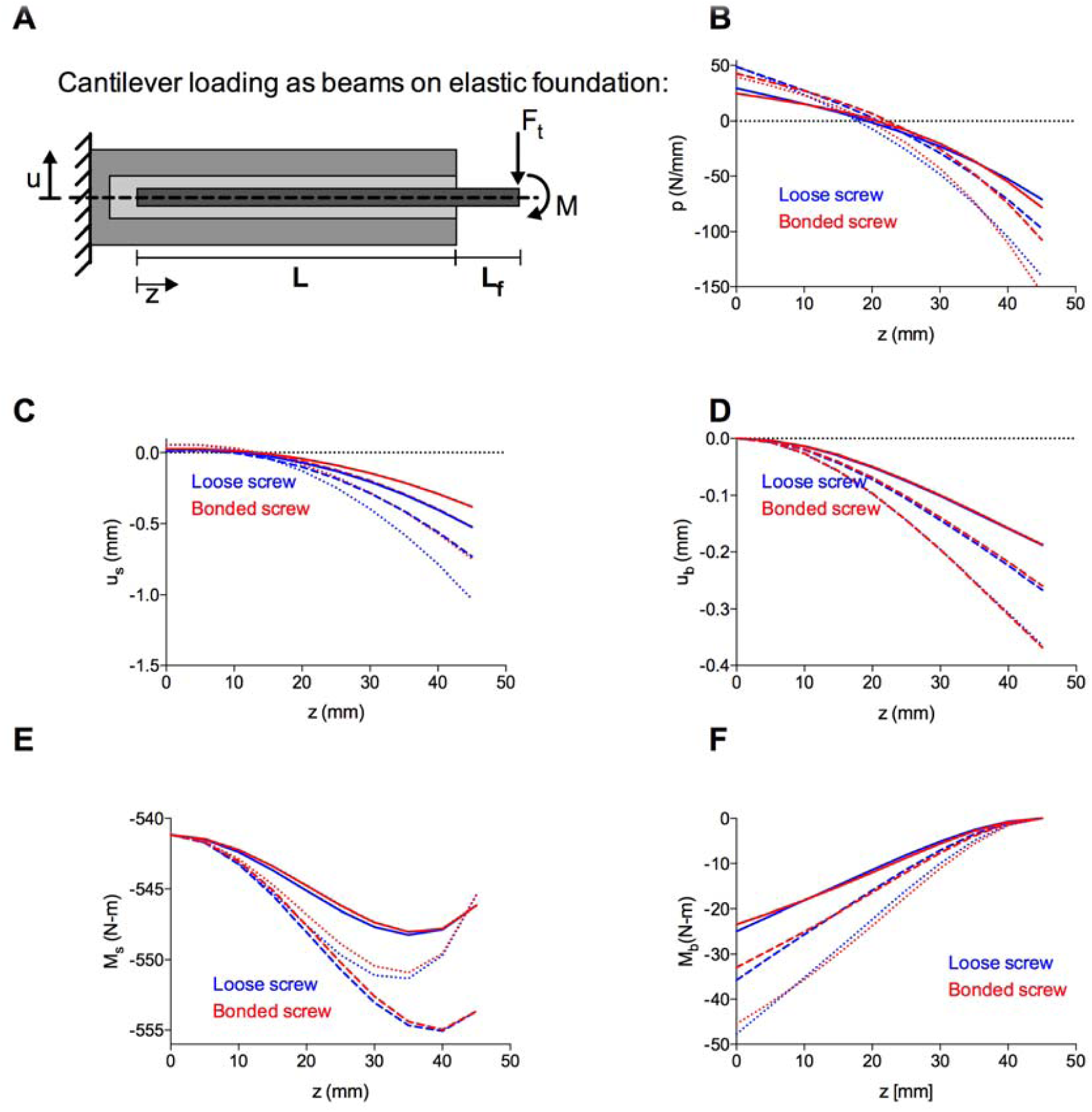
Load, displacement, and moment distribution along cantilevered screw. (A) Schematic of cantilever loading with pedicle screws and pedicle bone modeled as beams on elastic foundation. (B) Load distribution (C) deflection of the screw (D) deflection of the bone (E) moment of the screw and (F) moment of the bone along the length of the screw, L.

**Supplemental Figure 5.**
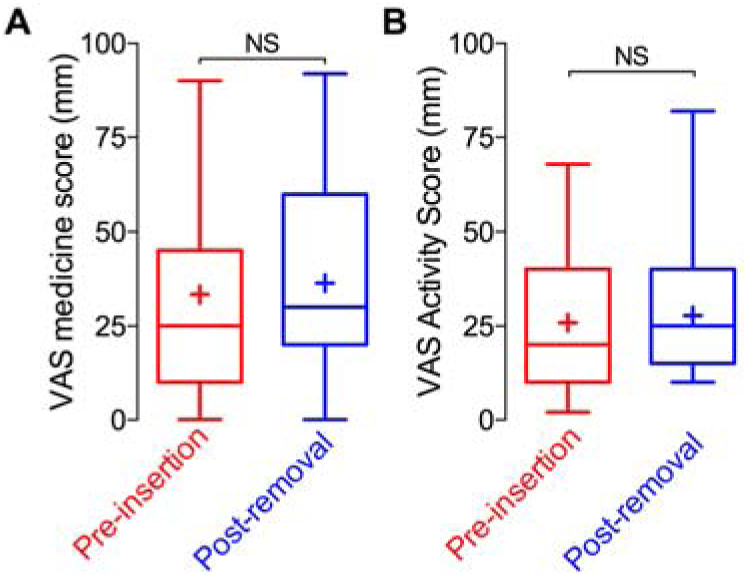
Analgesic use and activity VAS scores. (A) VAS analgesic medicine use assessment. (B) VAS physical activity assessment.

## APPENDICES

## Analytical stress modeling

A local stress analysis was considered to further assess pedicle screw loosening. Two cases of loading were considered: press fit loads caused by insertion of the screw into the pedicle and axial loading caused by force applied to the screw through the fixation system.

## Appendix 1 Cantilever screw loading

The transverse stiffness of the trabecular bone, C_t_, is given by

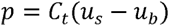

Where u_s_(z) and u_b_(z) are the deflections of the screw and the bone neutral axes, and p(z) is the continuously distributed transverse load that the trabecular bone exerts on the cortical bone and screw.

From beam theory,

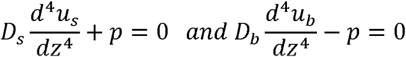

With D_s_ and D_b_ as the flexural stiffness of the screw and the bone (D=EI) which can be solved to give:

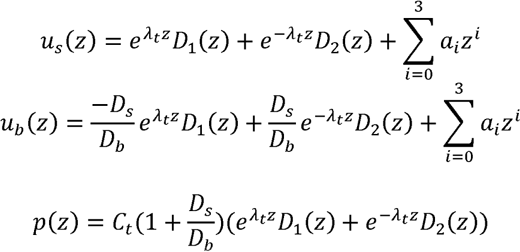

And

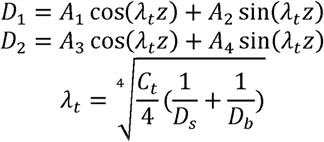

Where

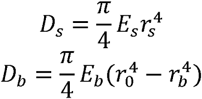

Using the boundary conditions:

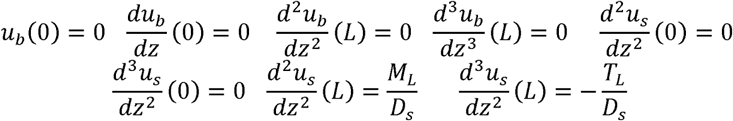

For example parameters F=100N, L=45mm, and M=0N-m the constants were found to be:

|  |  |  |  |
| --- | --- | --- | --- |
| $a_0$ | $1.96 \times 10^{-6}$ | $A_1$ | $-2.19 \times 10^{-6}$ |
| $a_1$ | $3.05 \times 10^{-4}$ | $A_2$ | $3.50 \times 10^{-6}$ |
| $a_2$ | $2.93 \times 10^{-2}$ | $A_3$ | $-3.30 \times 10^{-6}$ |
| $a_3$ | $6.10 \times 10^{-2}$ | $A_4$ | $3.51 \times 10^{-6}$ |

The bending moments of the screw and the bone can be found from:

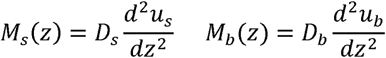

The stress resulting from axial loading of the pedicle screw can then be defined as

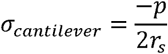

Representative graphs for the load distribution, screw and bone deflection, and moment distribution are shown in Supplementary Figure 4 for standing, flexion, and extension conditions.

The octahedral stress was found with σ_1_=σ_r_+σ_cantilever_ σ_2_=σ_θ_, and σ_3_=0.

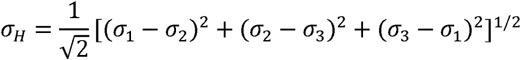

The octahedral stress was calculated as the pedicle screw diameter difference changed (x-axis) and plotted against octahedral stress (y-axis).

Pedicle screw diameter difference was defined as

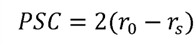

Pedicle screw diameter difference in the model was varied by changing the diameter of the pedicle.

**Table A.1.** Variables and assumed constants.

| Parameter |  |  |
| --- | --- | --- |
| $E_s$ | screw modulus | 180 GPa |
| $E_b$ | bone modulus | 21 GPa |
| $E_c$ | trabecular bone modulus | 42.8 MPa |
| $r_s$ | screw radius | 2.25-4.25 mm |
| $r_0$ | outer radius of cortical bone | 2.5-9.5 mm |
| $r_b$ | inner radius of cortical bone | 1.3-4.9 mm |
| $\rho$ | $r_s/r_0$ | |
| $\nu_{tb}$ | poisson ratio of trabecular bone | 0.37 |
| $\nu_{cb}$ | poisson ratio of cortical bone | 0.25 |
| $L$ | screw length | 35-60 mm |
| $L_f$ | screw head length | 10 mm |
| $F$ | transverse applied load | |
| $D_s$ | flexural stiffness of screw (EI) | |
| $D_b$ | flexural stiffness of bone (EI) | |
| $C_t$ | transverse stiffness of elastic trabecular bone | |
| $\alpha$ | rigidly connected vs. loose screw coefficient | |
| $p$ | continuously distributed transverse load | |

**Table A.2.** Physiologic loading conditions.

| Loading Conditions | Standing [1] | Flexion [2] | Extension [2] |
| --- | --- | --- | --- |
| F | 500N | 1175 N | 500 N |
| M | 0 N-m | -7.5 N-m | 7.5 N-m |

## Appendix 2 Screw Interference Fit

Under the assumptions of thick walled cylinders, the radial stresses are obtained from summing the forces for an element:

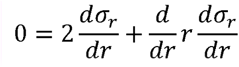

Integrating, the radial stress is defined as

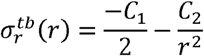

For small dθ, the circumferential stress is defined as

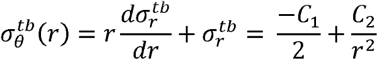

Applying the boundary conditions

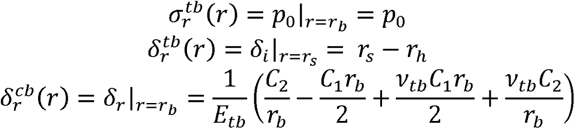

The following equations were used to find the coefficients C_1_, C_2_, and p_0_.

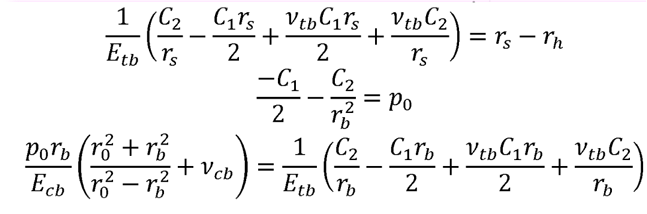

For the example parameters r_s_=4.5mm and r_0_=10mm, the constants were found to be:

|  |  |
| --- | --- |
| $C_1$ | $5.208 \times 10^6$ |
| $C_2$ | $1.086 \times 10^2$ |
| $p_0$ | $-4.124 \times 10^6$ |

Assuming that the screw modulus is much greater than the trabecular bone modulus (E_s_, >>E_tb_), we can assume E_s_ goes to infinity and that only the bone deforms: *δ_i_* = *δ_rh_* − *δ_rs_* = *r_s_*; − *r_h_*. Holes were tapped 1mm diameter smaller than the screw size such that *δ_i_* = *r_s_*; − *r_h_*= 0.5 mm, where *r_s_* is the radius of the pedicle screw, and *r_h_* is the radius of the tapped hole prior to screw insertion.

The stresses were then found at the screw-trabecular bone interface:

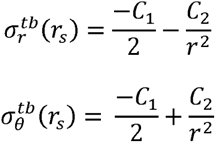

For the stress profile in the cortical bone, the solved equations for an internally pressurized thick-walled cylinder were used

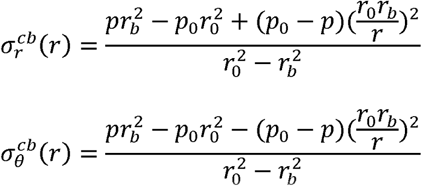

Where p_0_, the pressure outside the cortical bone, is zero, r_0_ is the radius of the cortical bone, r_b_ is the radius of the trabecular bone, and p is the internal pressure at the trabecular-cortical bone interface.

